# Human hematopoietic stem cell vulnerability to ferroptosis

**DOI:** 10.1101/2022.06.03.494357

**Authors:** Jiawei Zhao, Yuemeng Jia, Dilnar Mahmut, Amy A. Deik, Sarah Jeanfavre, Clary B. Clish, Vijay G. Sankaran

## Abstract

Hematopoietic stem cells (HSCs) have a number of unique physiologic adaptations that enable lifelong maintenance of blood cell production, including a highly regulated rate of protein synthesis. Yet the precise vulnerabilities that arise from such adaptations have not been fully characterized. Here, inspired by a bone marrow failure disorder due to loss of the histone deubiquitinase MYSM1, characterized by selectively disadvantaged HSCs, we show how reduced protein synthesis in HSCs results in increased ferroptosis. HSC maintenance can be fully rescued by blocking ferroptosis, despite no alteration in protein synthesis rates. Importantly, this selective vulnerability to ferroptosis not only underlies HSC loss in MYSM1 deficiency, but also characterizes a broader liability of human HSCs. Increasing protein synthesis rates via MYSM1 overexpression makes HSCs less susceptible to ferroptosis, more broadly illuminating the selective vulnerabilities that arise in somatic stem cell populations as a result of physiologic adaptations.

## Introduction

Lifelong production of blood and immune cells, or hematopoiesis, requires the continuous activity of rare hematopoietic stem cells (HSCs) that reside in the bone marrow (Laurenti and Göttgens, 2018; Liggett and Sankaran, 2020). These HSCs primarily remain in a quiescent state and can be activated to maintain hematopoiesis in homeostatic conditions or under the context of stressors, such as infections (Baldridge et al., 2010). Alterations to HSC function can cause bone marrow failure or promote progression to pre-malignant states and blood cancers. As a result of the sensitivity of HSCs to genotoxic or other stress-induced insults that can impair their effectiveness over the lifespan, these cells have acquired a variety of adaptations that can help to sustain their function. HSCs have a number of metabolic adaptations that include reliance on glycolysis and autophagy to enable sufficient energy generation without excess oxidant stress (Dong et al., 2021; Ho et al., 2017; Mortensen et al., 2011; Tothova et al., 2007). In addition, HSCs are characterized by low and highly-regulated rates of protein synthesis, which needs to be balanced with the rate of protein degradation, to maintain effective blood cell production (Kruta et al., 2021; Magee and Signer, 2021; Signer et al., 2014, 2016). While considerable insights have been generated from studies of HSCs conducted through targeting specific genes or pathways, the full spectrum of adaptations found in HSCs and the vulnerabilities arising as a result remain incompletely understood.

Studies of human disorders characterized by HSC loss or dysfunction provide a unique opportunity to gain new insights into the selective vulnerabilities found in these cells (Liggett and Sankaran, 2020). For example, studies of Fanconi anemia, which is characterized by a variety of mutations in the homologous recombination repair pathway, have revealed a role for these proteins in preventing HSC genotoxicity resulting from endogenous aldehydes (Garaycoechea et al., 2012, 2018). In an analogous manner, we reasoned that studies of a bone marrow failure disorder attributable to biallelic loss-of-function of the histone deubiquitinase MYSM1 could reveal additional and previously unappreciated pathways that are important for human HSC maintenance. The phenotype of these patients is distinct from other bone marrow failure syndromes and therefore this disease provides an opportunity to identify new regulatory aspects found in HSCs. In this disorder, there is progressive HSC loss with a reduced level of mature blood and immune cell production. Importantly, HSC transplantation and somatic reversion can rescue the phenotype observed in these patients (Alsultan et al., 2013; Belle et al., 2020; Le Guen et al., 2015; Li et al., 2020b; Zhan et al., 2021). Here, we set out to faithfully model this disorder with genome editing in human HSCs and conduct in depth mechanistic studies to decipher how this loss of MYSM1 can compromise HSC function. Our studies reveal a selective vulnerability of all human HSCs to ferroptosis and uncover a unique physiologic role for this distinct form of cell death in human hematopoiesis.

## Results

### Loss of MYSM1 compromises human HSC function

Bone marrow failure syndrome 4 (OMIM #618116) is a rare autosomal recessive genetic disorder characterized by mutations in histone deubiquitinase MYSM1. Over a dozen patients with this form of bone marrow failure due to a number of distinct mutations have been reported to date (Alsultan et al., 2013; Belle et al., 2020; Bluteau et al., 2018; Le Guen et al., 2015; Li et al., 2020b; Ulirsch et al., 2019; Zhan et al., 2021). The majority of these mutations cause complete loss-of-function of MYSM1 (Figure 1A). Importantly, this form of bone marrow failure is characterized by reduced production of all hematopoietic lineages and reduced cellularity of the bone marrow, suggesting a general defect impacting HSCs. Consistent with this, at least one patient has had somatic reversion in a subpopulation of HSCs enabling complete correction of their hematologic and immune phenotypes and other patients have been cured with HSC transplantation (Le Guen et al., 2015; Revy et al., 2019).

**Figure 1.**
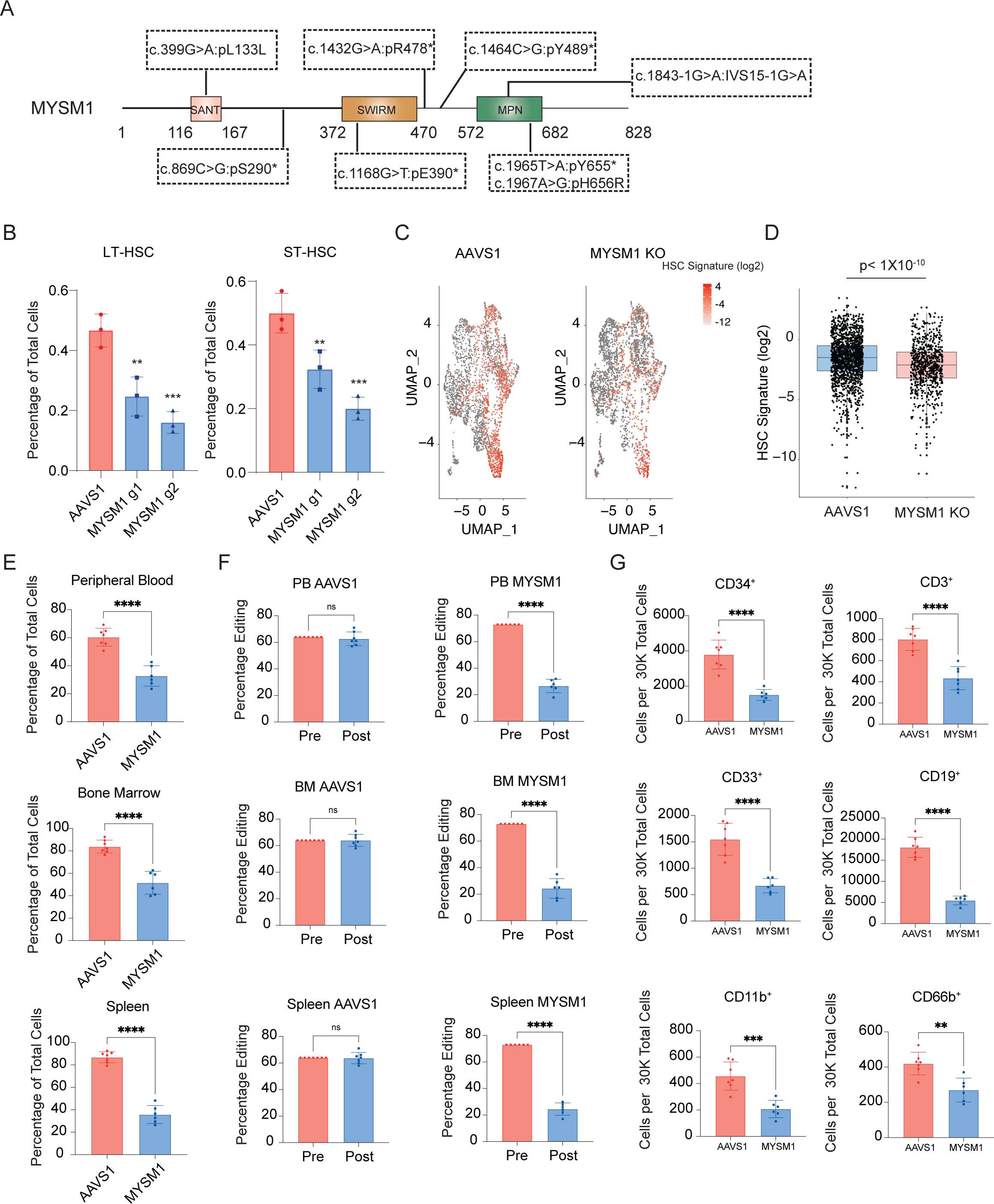
Loss of MYSM1 in CD34+ HSPCs recapitulates bone marrow failure phenotype observed in patients. (A) Comprehensive schematic diagram of MYSM1 mutants identified in patients. (B) Quantification of LT- and ST-HSC populations in AAVS1 and MYSM1 edited CD34^+^ HSPCs. (C) Uniform manifold approximation and projection (UMAP) plots of 4,293 (AAVS1) and 3,871(MYSM1 KO) CD34^+^CD45RA^-^CD90^+^ cells, highlighted according to average z-score normalized HSC gene signature. (D) Box plot of z-score normalized HSC signature expression of all cells of AAVS1 and MYSM1 KO groups. (E) Percentage of total human CD45+ cells after RBC depletion from indicated sites of NBSGW mice xenotransplanted with AAVS1 and MYSM1 edited cord blood CD34^+^ HSPCs. (F) Percentage of edits of cells before and after xenotransplantation from indicated sites of NBSGW mice xenotransplanted with AAVS1 and MYSM1 edited cord blood CD34+ HSPCs. (G) Cell numbers per 30,000 RBC depleted bone marrow cells of progenitor cells (CD34^+^), T cells (CD3^+^), myeloid (CD33^+,^ CD11b^+^), B cells (CD19^+^) and granulocytes (CD66b^+^).

While mouse knockouts of Mysm1 do display some hematologic phenotypes (Belle et al., 2015; Fiore et al., 2020; Huang et al., 2016; Huo et al., 2018; Won et al., 2014), there are notable differences in comparison to what is observed in human patients. To more fully define the human HSC defects arising with MYSM1 mutations, we sought to faithfully model the loss-of-function mutations observed in patients using CRISPR-Cas9 genome editing (Shen et al., 2021; Wagenblast et al., 2019). Two days after Cas9 ribonucleoprotein (RNP) delivery, we observed an overall editing efficiency of greater than 90% with two independent MYSM1 gRNAs (Figure S1A). Among the edits, the majority were predicted to result in nonsense mutations and a significant reduction of protein level was observed by three days post-editing (Figure S1B-C). Concomitantly, we observed a reduction of both long term (LT-) and short term phenotypic HSCs (ST-HSCs) (Figure 1B, S1D). Given the imprecision of surface markers to confidently delineate molecularly-defined HSCs (Liggett and Sankaran, 2020; Voit et al., 2022), we performed single cell RNA sequencing (scRNA-seq) on AAVS1 control and MYSM1 edited CD34^+^CD45RA^-^CD90^+^ cells and observed a significant reduction of cells harboring a molecular signature known to enrich for HSCs with MYSM1 editing (Bao et al., 2020) (Figure 1C-D).

To fully characterize the role of MYSM1 in human hematopoiesis, we transplanted AAVS1 control and MYSM1 edited cord blood CD34^+^ HSPCs into the NOD.Cg-Kit^W-41J^ Tyr ^+^ Prkdc^scid^ Il2rg^tm1Wjl^/ThomJ (NBSGW) strain of immunodeficient and Kit mutant mice (Fiorini et al., 2017; McIntosh et al., 2015; Voit et al., 2022). Consistent with the *in vitro* HSC depletion phenotype, there was significantly lower human CD45^+^ cell engraftment in the peripheral blood, bone marrow, and spleen at 4 months post-transplantation (Figure 1E, S1E), when hematopoiesis will predominantly be driven by transplanted HSCs. Importantly, the overall editing efficiency of cells was also dramatically reduced compared to CD34^+^ HSPCs before transplantation (Figure 1F), suggesting a strong selection against MYSM1 deficient cells. Based upon both the reduced bone marrow engraftment and depletion of MYSM1 edited alleles, we estimate an ∼8-fold loss of HSCs. Analysis of all major lineages confirmed that the repopulation defect broadly impacted all lineages (Figure 1G). Collectively, this data confirmed that normal MYSM1 expression is essential for human HSC maintenance both *in vitro* and *in vivo*, and genome editing of MYSM1 in HSPCs can faithfully recapitulate the bone marrow failure phenotypes observed in patients with biallelic mutations. Importantly, this model enabled us to ask critical questions about the molecular basis for HSC loss in this disorder, which remains enigmatic despite a number of phenotypic studies in mouse models.

### Reduced protein synthesis rates in HSCs with MYSM1 loss

To begin to define how MYSM1 loss can compromise human HSC function, we examined the scRNA-seq data for those cells that had the molecular HSC signature and found that there was a significant downregulation of gene sets and genes implicated in ribosomal biogenesis and translation regulation (Figure 2A-C). Given that this downregulation appeared to be subtle, we confirmed that the expression of various ribosomal proteins and eukaryotic initiation factor (EIF) encoding mRNAs was significantly reduced in CD34^+^CD45RA^-^CD90^+^ cells (Figure S2A). Moreover, gene set enrichment analysis (GSEA) reinforced the finding that genes involved in translation initiation and ribosome formation were significantly depleted in MYSM1 edited HSCs (Figure 2D, S2B, Table S1-2). Importantly, the protein levels of many ribosomal proteins were also significantly reduced (Figure 2E). These findings are consistent with a recent report using a Mysm1 knockout mouse, where Mysm1 was shown to be important for transcription of ribosomal protein genes through occupancy at the proximal promoters and transcriptional activation (Belle et al., 2020).

**Figure 2.**
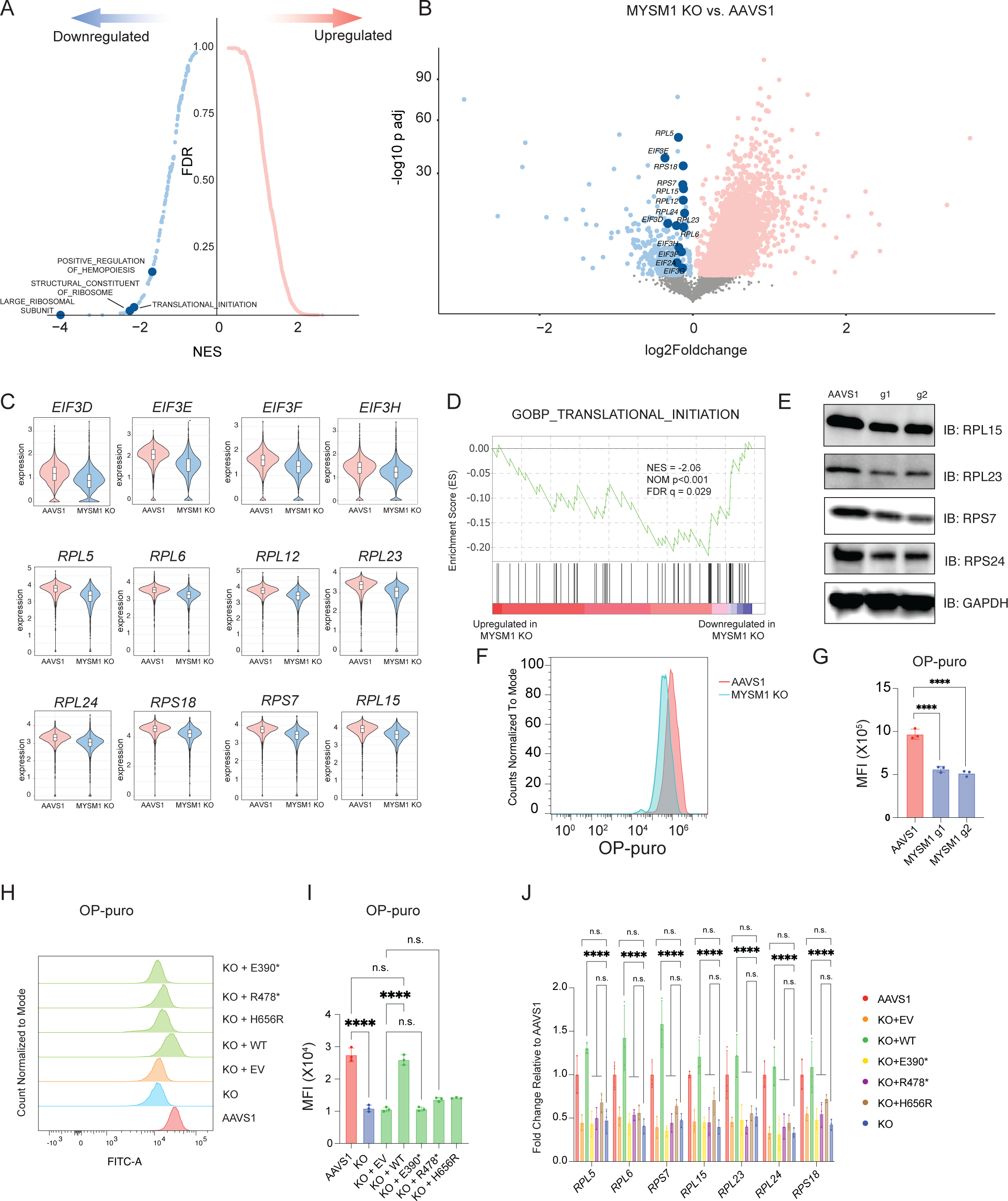
Loss of MYSM1 results in reduced cellular translation. (A) Top enriched signatures in upregulated and downregulated genes in MYSM1 KO cells with average z-score normalized HSC gene signature > 0.2, highlighting pathways directly related to translation regulation. (B) Volcano plot of MYSM1 KO vs. AAVS1 gene expression in HSCs, highlighting genes that are involved in ribosomal biogenesis and translational initiation. (C) Representative violin plots of ribosomal protein gene expression in AAV1 and MYSM1 KO HSCs. (D) Western blot analysis of representative ribosomal protein levels of CD34^+^CD45RA^-^CD90^+^ sorted cells with AAVS1 or MYSM1 editing. (E) Gene set enrichment analysis (GSEA) showing translation initiation is significantly downregulated in MYSM1 KO cells. (F) Representative flow cytometric histogram of O-Propargyl-puromycin based translation rate analysis on CD34^+^CD45RA^-^CD90^+^ sorted cells with AAVS1 or MYSM1 editing. (G) Quantification of mean fluorescence intensity (MFI) of O-Propargyl-puromycin based translation rate analysis on CD34^+^CD45RA^-^CD90^+^ sorted cells with AAVS1 or MYSM1 editing. (H) Representative flow cytometric histogram of O-Propargyl-puromycin based translation rate analysis on CD34^+^CD45RA^-^CD90^+^ sorted cells with AAVS1 or MYSM1 editing, and MYSM1 edited cells transduced with indicated overexpression constructs. (I) Quantification of mean fluorescence intensity (MFI) of O-Propargyl-puromycin based translation rate analysis on CD34^+^CD45RA^-^CD90^+^ sorted cells with AAVS1 or MYSM1 editing, and MYSM1 edited cells transduced with indicated overexpression constructs. (J) Real time PCR analysis of representative ribosomal protein expression of AAVS1 or MYSM1 edited cells, and MYSM1 edited cells transduced with indicated overexpression constructs.

While our data suggested that the global transcriptional program of ribosome biogenesis was downregulated, we sought to directly measure the impact on protein translation rates in HSCs. We therefore labeled CD34^+^CD45RA^-^CD90^+^ cells with O-Propargyl-puromycin (OP-puro) to assess translation rates in this population using flow cytometry (Hidalgo San Jose et al., 2020; Signer et al., 2014). The rate of protein translation rate was reduced to ∼50% in MYSM1 edited HSCs compared to AAVS1 control edited cells (Figure 2F-G), directly demonstrating altered mRNA translation due to the loss of MYSM1 that causes reduced transcription of ribosomal proteins and other genes (Belle et al., 2020). Importantly, we could confirm that this loss of protein synthesis was specifically due to MYSM1 activity by lentiviral rescue of edited cells, which could restore OP-puro incorporation and ribosomal protein gene expression to the levels observed in healthy HSCs (Figure 2H-J). Importantly, several mutations found in patients were unable to elicit a similar rescue, validating the loss-of-function due to these alleles (Figure 2H-J).

### Compromised HSC function in MYSM1 deficiency due to ferroptosis

While we and others have observed reduced ribosomal protein gene expression upon MYSM1 loss with accompanying reduced protein synthesis, how this would compromise HSC function remained unclear. HSCs are known to have a highly regulated and generally low rate of protein synthesis (Signer et al., 2014), but the specific liabilities that arise from these adaptations have not been fully defined. To identify additional causes for the HSC loss, we examined the gene expression changes identified through scRNA-seq. Interestingly, we observed a large number of upregulated genes in MYSM1 edited HSCs, suggesting that some of these genes may provide insights into altered pathways underlying the observed loss of HSCs. Among the upregulated pathways, we found that there was a significant enrichment in oxidant detoxification, antioxidant, and iron binding genes (Figure 3A, S3B, Table S1-2). Genes involved in iron metabolism and antioxidant activity were significantly upregulated, whereas genes involved in iron absorption, such as TFRC, were significantly reduced (Figure 3B-C, S3A). These data suggested that the cells might undergo significant oxidative stress and this may be accompanied by dysregulated iron handling.

**Figure 3.**
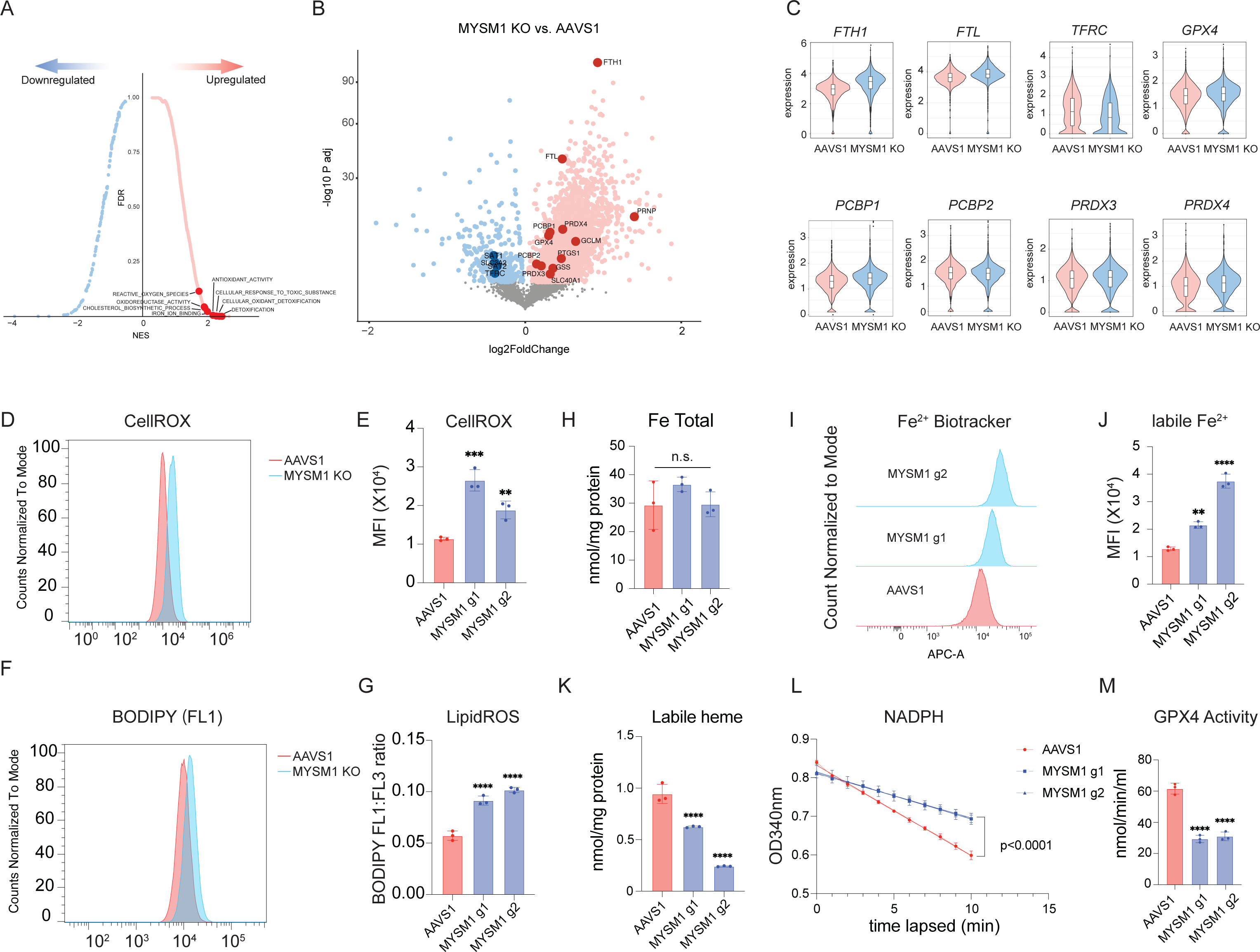
Loss of MYSM1 disrupts iron handling and results in increased ferroptosis in HSCs. (A) Top enriched signatures in upregulated and downregulated genes in MYSM1 KO cells with average z-score normalized HSC gene signature > 0.2, highlighting pathways relevant to iron metabolism and ferroptosis. (B) Volcano plot of MYSM1 KO vs. AAVS1 gene expression in HSCs, highlighting genes that are involved in iron transport, storage and metabolism. (C) Representative violin plots of iron transport, storage and metabolism gene expression in AAVS1 and MYSM1 KO HSCs. (D) Representative flow cytometric histogram of cellular ROS level of sorted CD34^+^CD45RA^-^CD90^+^ cells measured by CellROX dye. (E) Quantification of mean fluorescence intensity (MFI) of cellular ROS level of sorted CD34^+^CD45RA^-^CD90^+^ cells measured by CellROX dye. (F) Representative flow cytometric histogram of oxidized BODIPY dye of sorted CD34^+^CD45RA^-^CD90^+^ cells. (G) Quantification of cellular lipid peroxidation level of sorted CD34^+^CD45RA^-^CD90^+^ cells measured by ratio of oxidized and non-oxidized BODIPY dye. (H) Quantification of total intracellular iron level of AAVS1 and MYSM1 edited cells. (I) Representative flow cytometric histogram of ferrous iron level of sorted CD34^+^CD45RA^-^CD90^+^ cells as measured by Fe^2+^ biotracker dye. (J) Quantification of mean fluorescence intensity (MFI) of labile ferrous iron (Fe^2+^) level of sorted CD34^+^CD45RA^-^CD90^+^ cells as measured by Fe^2+^ biotracker dye. (K) Quantification of total intracellular labile hemin level of AAVS1 and MYSM1 edited cells. (L) Time lapsed reduction of NADPH availability in GPX4 assay reaction measure by OD340. (M) Calculated intracellular GPX4 activity of sorted CD34^+^CD45RA^-^CD90^+^ cells of AAVS1 and MYSM1 edited cells.

To directly assess how these gene expression changes were impacting the cells, we measured overall oxidative stress in CD34^+^CD45RA^-^CD90^+^ cells using the dye CellROX, which can measure all reactive oxygen species (ROS), and found that MYSM1 edited cells displayed globally increased ROS levels (Figure 3D-E). Moreover, we found that lipid peroxidation levels, as measured by staining with the lipid dye BODIPY, were significantly increased in MYSM1 edited HSCs (Figure 3F-G). While we saw gene expression changes in iron regulating genes that would suggest protection from a process such as ferroptosis (Dixon and Stockwell, 2014; Dixon et al., 2012; Stockwell et al., 2017), the elevated lipid peroxidation suggested increased ferroptosis in this setting. Indeed, while we observed no change of total intracellular iron levels between AAVS1 control and MYSM1 edited HSCs, the intracellular concentration of redox active ferrous iron (Fe^2+^) was significantly higher in MYSM1 edited HSCs than controls (Figure 3H-J). Interestingly, one of the key iron handling processes (Hentze et al., 2010), heme synthesis, appeared to be inhibited and intracellular labile heme levels were also reduced in MYSM1 edited cells (Figure 3K). Additionally, although we observed a transcriptional upregulation of *GPX4* mRNA in the HSC scRNA-seq data, the GPX4 activity in MYSM1 edited HSCs was significantly reduced (Figure 3L-M). Together, our data suggested that MYSM1 edited cells suffer from a significant upregulation of ROS that may underlie the HSC dysfunction and which appeared to arise from impaired iron handling, suggestive of ferroptosis being activated in this setting.

To further characterize the observed alterations that accompany the induction of ferroptosis in HSCs, we performed lipidomic analyses on the CD34^+^CD45RA^-^CD90^+^CD133^+^ HSC-enriched fraction of AAVS1 control and MYSM1 edited cells. We also compared the MYSM1 edited HSC enriched fraction to the non-HSC fraction of bulk progenitors that are CD34^+^CD90^-^. Consistent with prior analyses in the setting of ferroptosis (Yang et al., 2016; Zhang et al., 2019), there was a significant depletion of a wide spectrum of polyunsaturated phospholipid species, which are known to be most profoundly impacted with the lipid peroxidation that characterizes ferroptosis (Figure 4A-C, S4A-B, Table S3-5). Critically, these changes were only observed within the HSC enriched compartment upon MYSM1 editing and were absent from the more differentiated progenitor compartment (CD34+CD90-). Importantly, while previous studies have focused on ester linked phospholipids, our data shows that the vast majority of phospholipids depleted are ether linked phospholipids, specifically the 1-*O*-alkenyl-glycerophospholipids in human HSCs, whose functional importance in ferroptosis has been highlighted in recent studies (Zou et al., 2020) (Figure 4D-E, S4C-D, Table S3-5). Indeed, our data shows that while a few ester linked phospholipids are depleted, the majority of depleted lipids are ether linked phospholipids (Figure 4F). These data reveal that ether linked, but not ester linked, phospholipids are more susceptible to lipid peroxidation and ferroptosis due to loss of MYSM1 in HSCs. Accompanying these changes, we also noted an upregulation of triglyceride species, which may be due to changes following lipid peroxidation in the setting of ferroptosis (Figure 4F).

**Figure 4.**
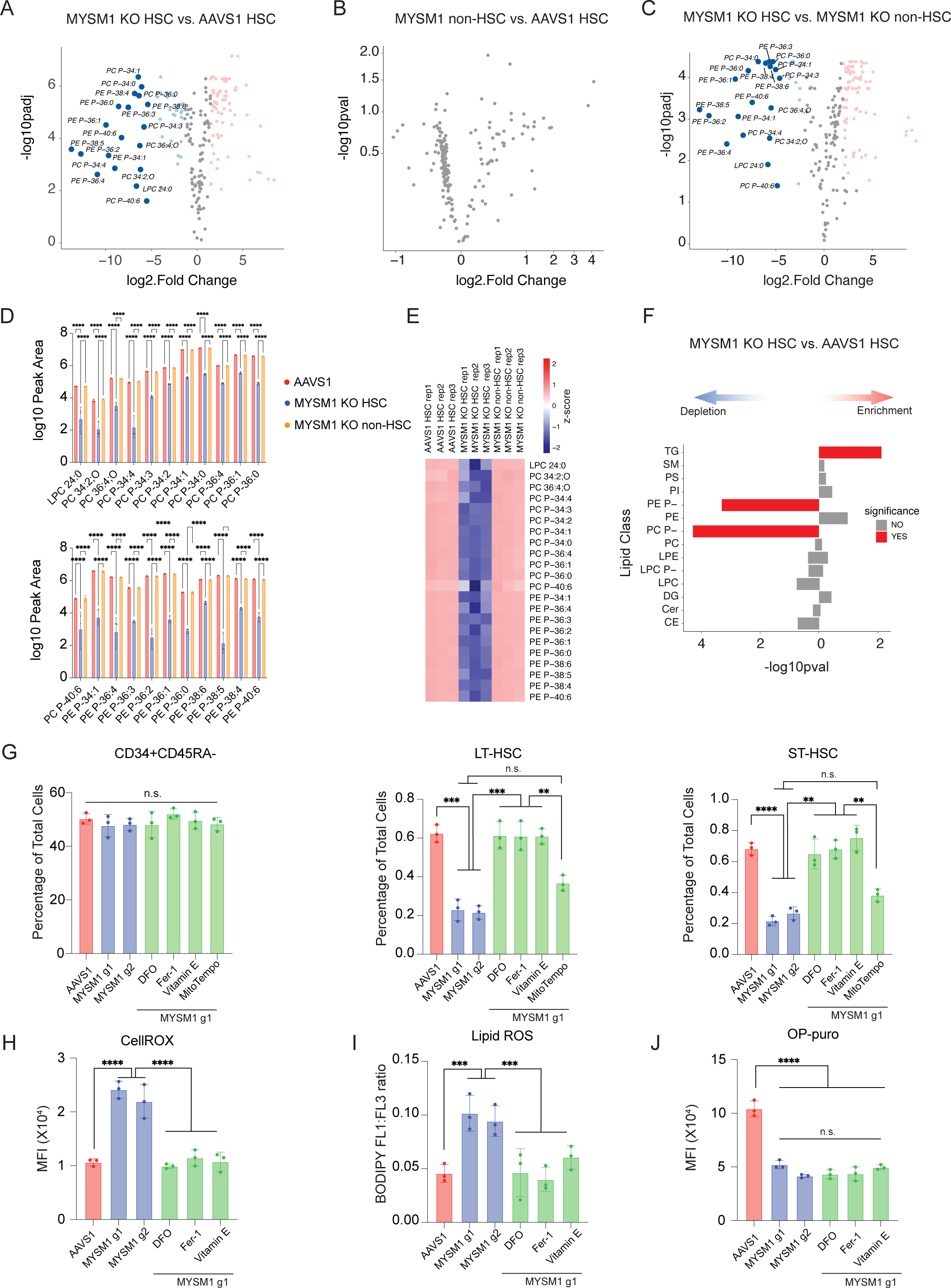
Inhibition of ferroptosis rescues HSC depletion due to MYSM1 loss. (A) Volcano plots of differentially expressed lipids of sorted CD90^+^CD133^+^ MYSM1 KO cells compared to AAVS1 control. (B) Volcano plots of differentially expressed lipids of sorted CD90^-^ MYSM1 KO cells compared to AAVS1 control. (C) Volcano plots of differentially expressed lipids of sorted CD90^+^CD133^+^ MYSM1 KO cells compared to CD90^-^ MYSM1 KO cells. (D) Representative bar plots of lipids depleted in CD90^+^CD133^+^ MYSM1 KO cells. (E) Heatmap of lipids depleted in CD90^+^CD133^+^ MYSM1 KO cells. (F) Lipid species enrichment of CD90^+^CD133^+^ MYSM1 KO cells compared to AAVS1 controls by over-representation analysis. TG: triacylglycerol; SM: sphingomyelin; PS: phosphatidylserine; PI: phosphotidylinositol; PE P-: 1-*O*-alkenyl-glycerophosphoethanolamine; PE: diacyl glycerophosphoethanolamine; PC P-: 1-*O*-alkenyl-glycerophosphocholine; PC: diacyl glycerophosphocholine; LPE: lyso-diacyl glycerophosphoethanolamine; LPC P-: lyso-1-*O*-alkenyl-glycerophosphocholine; DG: diacylglycerol; Cer: ceramide; CE: cholesterol ester. (G) Quantification of CD34^+^CD45RA^-^, ST-HSC, and LT-HSC populations in AAVS1 or MYSM1 edited cells, and MYSM1 edited cells treated with indicated chemicals. (H) Quantification of mean fluorescence intensity (MFI) of cellular ROS level of sorted CD34^+^CD45RA^-^CD90^+^ cells with AAVS1 or MYSM1 editing, and MYSM1 edited cells with indicated chemical treatments measured by CellROX dye. (I) Quantification of cellular lipid peroxidation level of sorted CD34^+^CD45RA^-^CD90^+^ cells with AAVS1 or MYSM1 editing, and MYSM1 edited cells with indicated chemical treatments measured by ratio of oxidized and non-oxidized BODIPY dye. (J) Quantification of mean fluorescence intensity (MFI) of O-Propargyl-puromycin based translation rate analysis on CD34^+^CD45RA^-^CD90^+^ sorted cells with AAVS1 or MYSM1 editing, and MYSM1 edited cells with indicated chemical treatments.

We next sought to directly test whether ferroptosis plays a role in the HSC depletion observed due to MYSM1 loss. We treated cells harboring loss of MYSM1 with a number of well established ferroptosis inhibitors including deferoxamine (DFO), ferrostatin-1 (Fer-1), and vitamin E, as well as the mitochondrially targeted antioxidant mitoTempo that will reduced ROS in the mitochondria, but will not alter ferroptosis (Li et al., 2020a). Critically, all ferroptosis inhibitors could fully rescue the loss of LT- and ST-HSCs due to MYSM1 deficiency without any alteration in the parental CD34^+^CD45RA^-^ progenitor population (Figure 4G). In contrast and consistent with the specificity of this rescue, the mitochondrially targeted antioxidant mitoTempo did not rescue the LT- or ST-HSC populations (Figure 4G). Moreover, all ferroptosis inhibitors were able to fully rescue and reduce the elevation of pan-ROS levels and lipid peroxidation in MYSM1 edited cells without impacting the compromised protein synthesis rates (Figure 4H-J, S4E). This finding shows how ferroptosis underlies the loss of HSCs in MYSM1 deficiency and lies downstream of the reduced protein synthesis that occurs in this disorder.

### Induction of ferroptosis in MYSM1 deficient HSCs due to translational downregulation of protective factors

Having identified increased ferroptosis as a vulnerability underlying dysfunction of HSCs with MYSM1 loss, we wanted to better delineate the underlying mechanisms. Though iron transport, storage, and metabolism as well as antioxidant genes were transcriptionally upregulated or remained unchanged, the levels of key proteins including GPX4, SLC7A11, and FTH1 were all significantly reduced in HSPCs and most profoundly reduced in the primitive HSC-enriched CD34^+^CD45RA^-^CD90^+^ compartment (Figure 5A-B). Given the observed reduction in global protein synthesis, we wondered if this downregulation could specifically impact certain pathways more selectively, including the factors that would ordinarily constrain ferroptosis, analogous to our prior findings in human hematopoietic lineage commitment (Khajuria et al., 2018). To directly assess this and define specific alterations in mRNA translation in an unbiased manner, we performed ribosome profiling with massively parallel sequencing (ribo-seq) to analyze specific changes in mRNA translation upon loss of MYSM1 (Ingolia et al., 2019). Quality control using triplet periodicity and other features confirmed the production of high-quality ribo-seq data from these rare primary cell populations (Figure S5A). Importantly, we observed MYSM1 as one of the most significantly downregulated genes (Figure 5C), which is expected given the loss-of-function edits that occur in the gene and serves to validate the assay. Strikingly, we found that genes involved in protection from ferroptosis were significantly downregulated in terms of translational efficiency (TE) in MYSM1 edited cells, in contrast to the observed transcriptional upregulation of the mRNAs encoding these genes. Pathways that were transcriptionally enriched such as ferroptosis, cholesterol and lipid synthesis, and antioxidant activity were all ranked among the most depleted pathways (Figure 5D-E, S5B, Table S6-7). Moreover, analysis of the TEs for specific genes confirmed a significant reduction. In contrast, pathways that were transcriptionally depleted were highly enriched pathways from the TE analysis (Figure 5D, S5C), suggesting a reciprocal compensatory mechanism between transcription and translation across many pathways in MYSM1 edited HSCs. A motif analysis on 5’UTRs of translationally downregulated mRNAs identified 5’UTR motifs that tend to be translated less efficiently due to loss of MYSM1 and demonstrated that these motifs were enriched on factors that ordinarily serve to constrain ferroptosis, providing a molecular basis for the observed selective downregulation of these factors (Figure 5G-H). Together, our data showed that reduced translation rates due to loss of MYSM1 result in HSC depletion through less efficient translation of ferroptosis protective genes.

**Figure 5.**
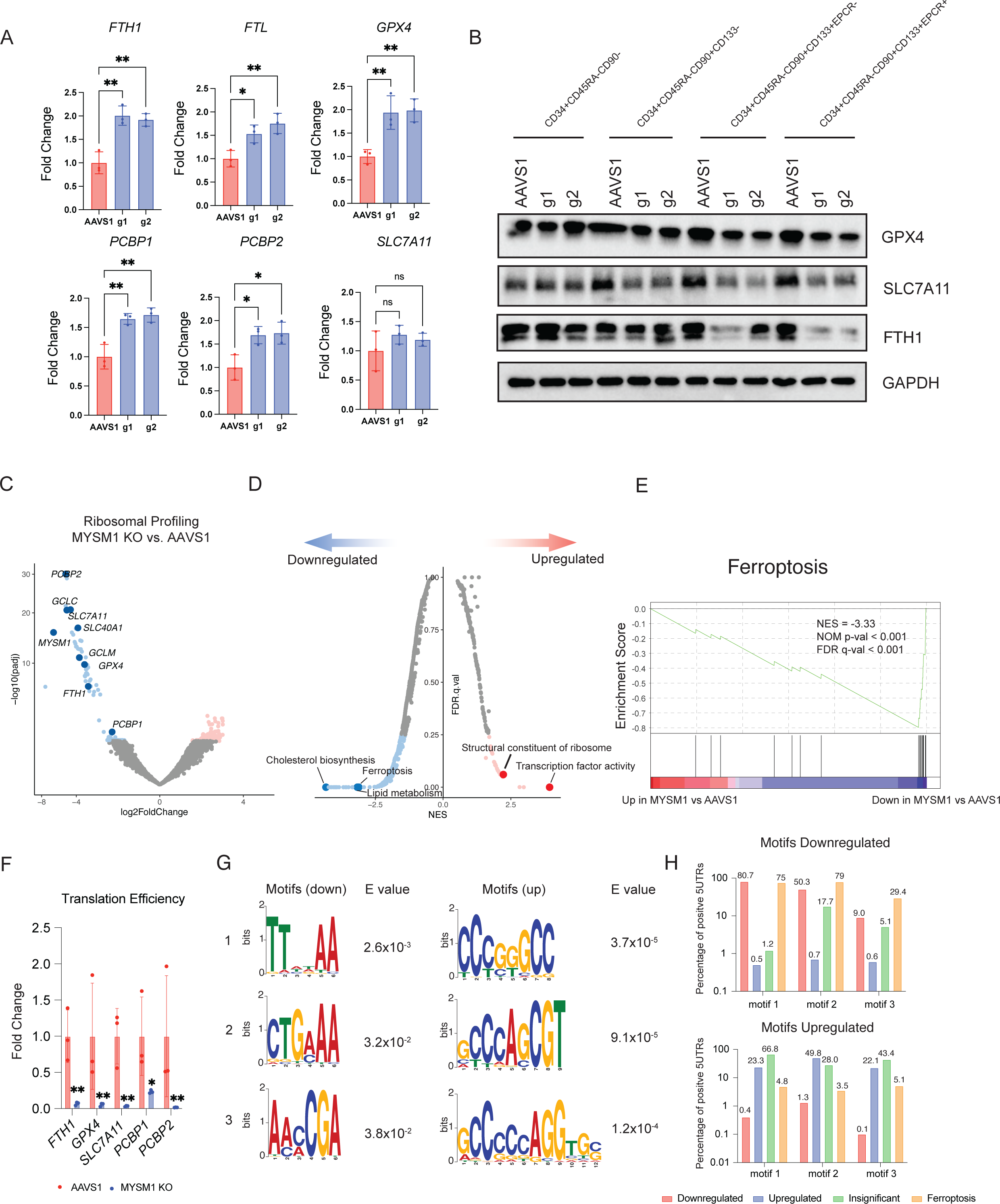
Reduced cellular translation due to loss of MYSM1 preferentially decreases translation efficiency of iron transport, storage and metabolism genes, resulting in overall decrease of protein level. (A) Real time PCR analysis of iron transport, storage and metabolism gene expression of sorted CD34^+^CD45RA^-^CD90^+^CD133^+^ cells. (B) Western blot analysis of key iron handling and ferroptosis genes in indicated fractions of culture CD34^+^ HSPCs edited by AAVS1 control or MYSM1. (C) Volcano plot of MYSM1 KO vs. AAVS1 differentially translated genes, highlighting genes directly relevant to cellular iron transport, storage and metabolism, and ferroptosis. (D) Top enriched signatures in differentially translated genes in MYSM1 KO cells. (E) Gene set enrichment analysis (GSEA) showing ferroptosis protective gene translation is significantly impaired in MYSM1 KO cells. (F) Representative translation efficiency change of ferroptosis genes in MYSM1 KO cells. (G) Motif discovery analysis identified genes with specific 5’ UTR motif pattern tend to be less or more efficiently translated due to MYSM1 loss. (H) Percentage of 5’UTRs of genes of different groups in downregulated or upregulated motifs discovered in (G)

### Healthy HSCs are selectively vulnerable to ferroptosis due to a low level of protein synthesis

We have thus far focused on the compromised HSC function due to MYSM1 deficiency causing reduced protein synthesis and increased ferroptosis, as a result. Given that low and highly-regulated protein synthesis is a critical adaptation in healthy HSCs (Magee and Signer, 2021; Signer et al., 2014), we wondered if this vulnerability to ferroptosis may be more broadly present in HSCs. We employed several well-characterized small molecule inducers of ferroptosis including erastin, FIN56, FINO2, and RSL3, as well as genome editing of *GPX4* to perturb human HSPCs. Remarkably, in contrast to the bulk population of human HSPCs, healthy HSCs were selectively vulnerable to ferroptosis via all of these distinct and complementary approaches (Figure 6A-B, S6A). All of these perturbations that resulted in ferroptosis in HSCs, increased global ROS levels and lipid peroxidation in the primitive HSC-enriched CD34^+^CD45RA^-^CD90^+^ compartment (Figure 6C-D), reinforcing the notion that HSC loss in the setting of these perturbations is accompanied by the expected alterations that characterize ferroptosis.

**Figure 6.**
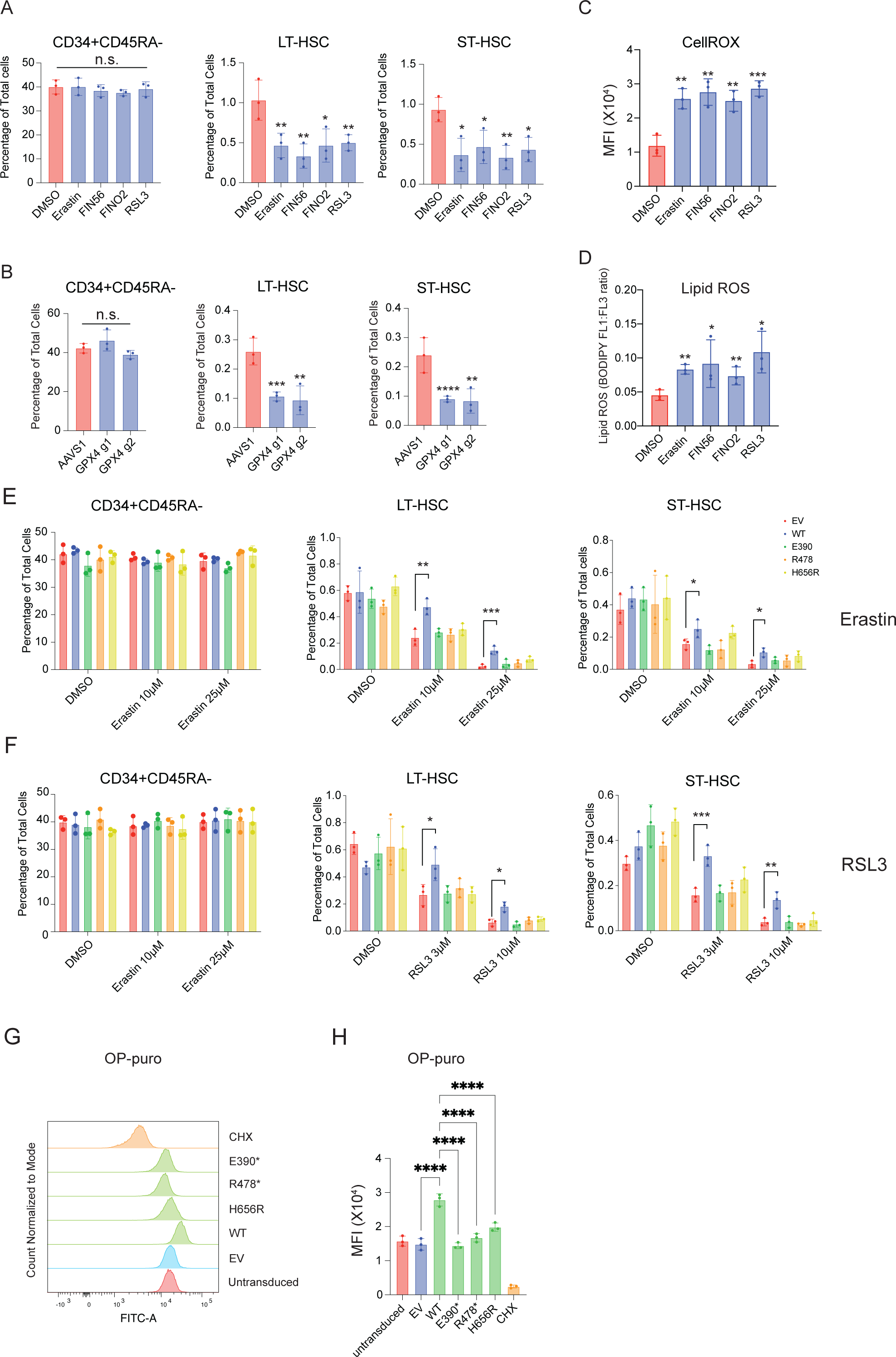
Normal hematopoietic stem cells (HSCs) are susceptible to ferroptosis induction and overexpression of MYSM1 can be protective due to increased translation. (A) Quantification of CD34^+^CD45RA^-^, ST-HSC, and LT-HSC populations in DMSO and indicated ferroptosis inducer-treated CD34^+^ HSPCs. (B) Quantification of CD34^+^CD45RA^-^, ST-HSC, and LT-HSC populations in AAVS1 or GPX4 edited cells. (C) Quantification of mean fluorescence intensity (MFI) of cellular ROS level of sorted CD34^+^CD45RA^-^CD90^+^ cells with DMSO or indicated ferroptosis inducer treatment measured by CellROX dye. (D) Quantification of cellular lipid peroxidation level of sorted CD34^+^CD45RA^-^CD90^+^ cells with DMSO or indicated ferroptosis inducer treatment measured by ratio of oxidized and non-oxidized BODIPY dye. (E) Quantification of CD34^+^CD45RA^-^, ST-HSC, and LT-HSC populations in DMSO and Erastin treated cells, and cells treated with Erastin and transduced with indicated constructs. (F) Quantification of CD34^+^CD45RA^-^, ST-HSC, and LT-HSC populations in DMSO and RSL3 treated cells, and cells treated with RSL3 and transduced with indicated constructs. (G) Representative flow cytometric histogram of O-Propargyl-puromycin based translation rate analysis on CD34^+^CD45RA^-^CD90^+^ sorted cells transduced with indicated constructs. (H) Quantification of mean fluorescence intensity (MFI) of O-Propargyl-puromycin based translation rate analysis on CD34^+^CD45RA^-^CD90^+^ sorted cells transduced with indicated constructs

Given that prior studies have shown that HSCs have a characteristically low level of protein synthesis (Signer et al., 2014) and we have demonstrated how reduced protein synthesis due to MYSM1 loss can induce ferroptosis, we wondered if overexpression of MYSM1 could augment protein synthesis in healthy HSCs and thereby protect the cells from ferroptosis. Induction of ferroptosis in the HSC compartment by either erastin or RSL3 at two different concentrations could be prevented by increasing the expression of normal MYSM1 (Figure 6E-F, S6B). Several disease-causal mutations in MYSM1 were unable to rescue this phenotype, reinforcing the need for full MYSM1 function to enable this rescue. Concomitantly, we observed an increase in protein synthesis rates in the HSC compartment with the normal but not disease-associated mutant forms of MYSM1 (Figure 6G-H). Collectively, we demonstrate how the low levels of protein synthesis that characterize even healthy HSCs confer a vulnerability to ferroptosis, which can be mitigated by augmenting protein synthesis in HSCs.

## Discussion

A milestone in our understanding of how the metabolic state of cells can be linked to cell survival and induction of cell death occurred ten years ago with the first description of ferroptosis (Dixon et al., 2012; Jiang et al., 2021; Stockwell et al., 2017). Since that time, a considerable number of studies have characterized the molecular regulation of this process (Bersuker et al., 2019; Ingold et al., 2018; Jiang et al., 2021; Yang et al., 2014), as well as how specific pathologic contexts can enable liabilities to cell death via ferroptosis, as exemplified by the sensitivity of cancer cells to ferroptosis (Hangauer et al., 2017; Jiang et al., 2015; Ubellacker et al., 2020; Viswanathan et al., 2017; Zou et al., 2019, 2020). However, despite the considerable advances made, the precise vulnerabilities of cell populations in physiologic settings to death via ferroptosis remains undefined. Here, through studies inspired by a rare inborn error impacting human HSCs due to deficiency of the histone deubiquitinase MYSM1, we have identified a distinct vulnerability of HSCs to ferroptosis. This sensitivity arises, at least in part, due to the low and highly-regulated rate of protein synthesis in these cells, which is thought to be critical for enabling homeostatic protein levels in these cells (Kruta et al., 2021; Signer et al., 2014). Through our in depth mechanistic studies, we are able to identify a key and previously undescribed connection that couples mRNA translation to the induction of ferroptosis and show how downregulation of mRNA translation impacts the translation of factors critical for ferroptosis, which harbor unique motifs in their 5’ UTRs. This adds to the emerging connections between the regulation of translation of specific factors, as is seen with selenoproteins, and sensitivity to ferroptosis (Alborzinia et al., 2022; Li et al., 2022).

While a considerable body of work has focused upon the metabolic adaptations that are found in HSCs and which enable unique adaptations, including the use of autophagy by these cells to regulate metabolism (Dong et al., 2021; Filippi and Ghaffari, 2019; Ho et al., 2017), the requirement for specific metabolites (Agathocleous et al., 2017; Cimmino et al., 2017), the fine-tuned regulation of iron homeostasis (Kao et al., 2018), and the need for proteostasis (Kruta et al., 2021; Mbong et al., 2018; Signer et al., 2014), the precise vulnerabilities arising from these adaptations remain poorly characterized. Human diseases that impact specific cell populations, as exemplified by MYSM1 deficiency that we study here, provide an opportunity to uncover the sensitivities of specific cells to activation of distinct pathways. In this context, we uncover a sensitivity of human HSCs to ferroptosis, which has broader implications in regulating these cells beyond the context of the rare bone marrow failure disorder that inspired this set of studies.

Beyond the insights gained into fundamental pathways critical in the regulation of human HSCs, our findings also have important translational and clinical implications. First, agents that can chelate intracellular iron may enable expansion of HSCs, as suggested through studies of eltrombopag for the treatment of aplastic anemia (Desmond et al., 2014; Kao et al., 2018; Olnes et al., 2012; Peffault de Latour et al., 2022), a condition characterized by loss of HSCs in the bone marrow. While some of the observed activities may be attributable to stimulation of the thrombopoietin signaling axis by eltrombopag, experimental studies do suggest a role for limiting intracellular iron in promoting improved maintenance and expansion of HSCs (Kao et al., 2018). Our findings emphasize the opportunities that may exist for helping preserve and augment HSC function in a number of bone marrow failure syndromes by restricting ferroptosis, including rare conditions such as MYSM1 deficiency, but also more common conditions characterized by HSC loss such as aplastic anemia and the myelodysplastic syndromes. Second, the observation that supraphysiologic oxygen can cause stress and impair HSC function may be attributable to the induction of lipid peroxidation and ferroptosis (Mantel et al., 2015). Blockage of ferroptosis may represent an ideal strategy for helping preserve HSC function in the setting of cell therapies requiring *ex vivo* manipulation. Third, ferroptosis is being actively pursued as a promising avenue for anti-cancer therapies, given the sensitivity of cancer cells to the induction of ferroptosis (Hangauer et al., 2017; Viswanathan et al., 2017). However, it is important to be aware of potential side effects of such agents and our findings suggest that HSC loss and resultant impaired hematopoiesis will be an issue that will require close monitoring as such agents are developed and tested. By further defining the underlying metabolic regulation that confers the observed vulnerability of HSCs to ferroptosis, a variety of clinical opportunities will likely arise to enable better treatments for therapy-resistant cancers and for intractable blood diseases.

## Acknowledgements

We thank members of the Sankaran lab and a number of colleagues, including P. van Galen, S. Schreiber, and B. Will, for valuable comments and advice on this work. This work was supported by the New York Stem Cell Foundation (NYSCF), a gift from the Lodish Family to Boston Children’s Hospital, the Edward P. Evans Foundation, and National Institute of Health (NIH) grants R01 DK103794 and R01 HL146500. V.G.S. is a NYSCF-Robertson Investigator.

## Author Contributions

J.Z. and V.G.S. conceived and designed the study. J.Z., D.M., and A.D. performed experiments. J.Z., Y.J., A.D., C.B.C., and V.G.S. analyzed data. J.Z. and V.G.S. wrote the paper with input from all authors. V.G.S. provided project oversight.

## Methods

### CD34+ HSPCs culture and nucleofection

Human CD34^+^ HSPCs were obtained from the Cooperative Center of Excellence in Hematology at the Fred Hutchinson Cancer Research Center. The cells were thawed and cultured in StemSpan SFEM II culture medium, supplemented with 1X CC100, recombinant thrombopoietin (TPO), and UM171. The RNP complex was assembled by mixing 50 pmol sgRNA and 75 pmol Cas9 protein and incubating at room temperature for 10-30 mins. The assembled RNP complex was delivered into CD34^+^ HSPCs by nucleofection using the Lonza 4D nucleofector system. The cells were harvested and genomic DNA was extracted 72 hours post-nucleofection. DNA fragments flanking the editing site at least 250 bp upstream and downstream were amplified and sent for Sanger sequencing to assess editing efficiencies. The .ab1 file was used for ICE analysis from Synthego website for prediction of editing efficiency and edit compositions (https://ice.synthego.com/#/).

### MYSM1 overexpression cloning and lentivirus packaging

*MYSM1* cDNA was amplified using CD34^+^ HSPC cDNA library and cloned into the pCRII vector using Zero Blunt TOPO cloning kit (Invitrogen). The sequence of amplified *MYSM1* cDNA was verified by Sanger sequencing before subcloning into the pLVX overexpression vector using the In-Fusion cloning kit (Takara). Q5 site directed mutagenesis kit was used to make H656R mutant.

For lentivirus packaging HEK-293T cells were cultured in DMEM medium supplemented with 10% FBS and 1X penicillin/streptomycin. After one passage, the cells were plated into 10 cm dishes at 50% confluency. On the next day (within 24 hrs after plating), WT and mutant MYSM1 transfer plasmids were co-transfected with VSV-G and psPAX2 plasmids at 1:1:1 molar ratio using linear PEI. Viral supernatants were harvested twice at 48 and 72 hours post-transfection. Combined viral supernatant was centrifuged and filtered at 0.45 μm. Filtered viral supernatant was then concentrated by ultracentrifugation with a 2 ml 20% sucrose cushion at 25,000 rpm in the SW32 rotor of Beckman Coulter ultracentrifuge. After ultracentrifugation, the supernatant was decanted and viral pellets were resuspended in ice cold PBS.

Concentrated virus was added to CD34^+^ HSPCs at an MOI of 50 in presence of 8 μg/ml polybrene. The cells were then spin infected at 2,000 rpm for 90 mins at 37C. After spin infection, the supernatant was removed and fresh culture medium was added. Puromycin selected was started 36 hours post-transduction.

### Xenotransplantation and animal model

All animal procedures were performed under the protocol approved by Boston Children’s Hospital Institutional Animal Care and Use Committee (IACUC). Cord blood was acquired from Dana Farber Cancer Institute’s Pasquarello tissue core. CD34^+^ HSPCs were enriched by EasySep Human Cord Blood CD34^+^ positive selection kit (StemCell Technologies). Enriched cells were nucleofected by AAVS1 control and MYSM1 sgRNAs. Three days post-nucleofection, a small fraction of cells will be harvested for ICE analysis of editing efficiency. The remaining cells will be intravenously injected into NBSGW immunodeficient mice (JAX#026622) at1 × 10^5^cells per mouse. Autoclaved sulfatrim antibiotic water was given and changed weekly to prevent potential infections. Peripheral blood, bone marrow, and spleen were harvested four months after transplantation for analysis of engraftment, MYSM1 editing efficiency and lineage-specific markers. Specifically, peripheral blood was harvested through cardiac puncture right after the mice were sacrificed and collected in EDTA coated tubes. Red blood cells were depleted by adding 20X volume of ddH_2_O for 30 seconds, the osmolarity was balanced by adding 10X PBS solution. If the pellet appears to be red, a second round of RBC lysis with the same method was used to deplete remaining RBCs. For bone marrow cell harvest, femurs and tibia were dissected and crushed by mortar and pestle in RPMI 1640 medium supplemented with 10% FBS. The cells were passed through 40 μm cell strainer. The cells were washed by ice-cold PBS once and washed cells were frozen down in 90% FBS + 10% DMSO. The majority of RBCs were automatically depleted by cryopreservation. For splenocyte isolation, spleens were dissected and minced. The minced spleen was digested by 1 mg/ml collagenase IV for 30 minutes at 37 degrees C. The digestion was stopped by addition of EDTA at a final concentration of 1 mM. Splenocytes were frozen in 90% FBS + 10% DMSO to deplete RBCs and stored in liquid nitrogen for subsequent analysis.

### Single cell RNA-seq analysis

Edited AAVS1 and MYSM1 KO cells were FACS sorted for CD34^+^CD45RA^-^CD90^+^ cells. The cells were resuspended at 1,000/μl. The scRNA-seq library was prepared using 10X Genomics scRNA-seq library prep kit according to manufacturer’s instructions. The sequencing results were pre-analyzed by the CellRanger pipeline to generate the matrix file, which was brought to downstream analysis. Normalization, scaling, and cell clustering was done in R by the Seurat v4.0.2 package. The top 1000 most highly variable genes were used to calculate the top 50 principal components. The HSC cluster was highlighted by averaging the z-score normalized expression of *CD34, HLF, CRHBP* for each cell, as previously described (Bao et al., 2020). The HSC population, defined by the averaged z score of more than 0.5 for the HSC signature, was used for DEG calling by DESeq2. The DEG list was used to generate rnk file, which was used to run the GSEA analysis (https://www.gsea-msigdb.org/gsea/index.jsp).

### Real time PCR analysis

The total RNA of sorted or unsorted cells were harvested using RNeasy mini RNA purification kit according to manufacturer’s instructions. 500 ng (sorted cells) to 1 μg (unsorted cells) of total RNA was used for reverse transcription using iScript cDNA synthesis kit (Biorad). The cDNA product was diluted at 1:20 and 2 μl of diluted cDNA was used for real time PCR analysis using iQ SYBR green supermix (Biorad). Data was normalized by loading control and presented as fold change compared to control samples.

### Immunoblotting analysis

Total protein lysate of sorted or unsorted cells was extracted by RIPA buffer in presence of protease inhibitor cocktail and PMSF on ice for 10 minutes followed by a 5-minute incubation with MNase at 37 C. The total lysate was then linearized by 1X SDS loading buffer and heated at 55 degrees C for 10 minutes. The lysate was run on 5-20% gradient gel and then transferred onto a PVDF membrane. The membrane was then blocked by 5% nonfat dry milk in TBS/T. The membrane was then incubated with primary antibodies at 1:1000 dilution at 4C overnight. HRP conjugated secondary antibodies were then incubated for 1hr at room temperature. Membrane was developed using ECL and imaged in the Biorad gel imaging system.

### Flow cytometric analysis

Cultured or freshly isolated cells were washed once by ice cold PBS, followed by staining of corresponding antibody cocktails at 1:60 dilution in FACS buffer (PBS+2%FBS) on ice for 20 minutes. The cells were then washed by FACS buffer twice and resuspended in FACS buffer and analyzed by BD LSRII and/or Fortessa flow cytometer.

### Intracellular ROS analysis

Nucleofected cells were sorted for CD34^+^CD45RA^-^CD90^+^ and stained with CellROX dye according to manufacturer’s instructions. Labeled cells were washed with ice cold PBS and analyzed by BD Accuri C6 table top flow cytometer.

### Lipid peroxidation analysis

Sorted cells were stained with BODIPY C11 lipid probe (Invitrogen) according to the manufacturer’s instructions. Labeled cells were washed and analyzed on BD Accuri C6 table top flow cytometer. For lipid peroxidation analysis, the peroxidation state of each group was calculated by mean fluorescence intensity of FL1 channel to that of FL3 channel.

### Cellular translation rate analysis

The cellular translation rate of sorted cells was measured by O-Propargyl-puromycin based translation assay kit (Cayman chemicals) according to manufacturer’s instructions. Briefly, the cells were stained with O-Propargyl-puromycin at a 1:40 dilution from the stock solution in culture medium at 37 degrees C for 30 minutes. Labeled cells were then washed with FACS buffer. The cells were fixed by the fixative reagent of the kit for 5 minutes at room temperature. Fixed cells were then stained with 5-FAM azide solution for 30 minutes at room temperature, which was then analyzed by flow cytometry.

### Total and ferrous iron measurement

Total cellular iron level was measured using iron colorimetric assay kit (BioVision) according to manufacturer’s instructions. Briefly, cells were harvested and homogenized in the iron assay buffer. Lysed cells were cleared by centrifugation at 15,000g for 5 minutes at 4 degrees C. The supernatant was collected and a small fraction was split for Qubit protein quantification. The rest of cell lysates were quantified for total iron after the reaction as a measurement of OD593. Iron concentrations were calculated by normalizing total protein concentration.

Intracellular ferrous iron was measured by labeling cells with BioTracker Far-red Labile Fe2+ Dye (Millipore Sigma) and analyzing by flow cytometry.

### Labile hemin analysis

Intracellular hemin level was measured by hemin colorimetric assay kit (BioVision) according to manufacturer’s instructions. Briefly, cells were lysed in the hemin assay buffer. 50 μl of cell lysates was used for quantification in 96 well plates. After a 30 minutes reaction, the plates were read for OD570. The final hemin concentration was normalized by total protein concentration.

### GPX4 activity analysis

The cellular GPX4 activity was measured indirectly by GPX4 inhibitor screening assay kit (Cayman Chemicals) according to manufacturer’s instructions. Briefly, collected cells were lysed in GPX4 sample buffer containing 50mM Tris-HCl, pH7.5, 5mM EDTA and 1mM DTT. 20 μl of sample was used for each reaction. Dynamic plate reading for OD340 was started immediately after all reaction components were added. Plates were read every 1 minute for a total 15 minutes. GPX4 activity was calculated according to the manufacturer’s user manual.

### Ribo-seq library preparation and analysis

Ribosome profiling sequencing libraries were constructed as previously described (Ingolia et al., 2012). Briefly, cells were treated with cycloheximide at final concentration of 100 μg/ml for 5 minutes at 37 degrees C. The cells were then harvested in presence of same concentration of cycloheximide to extract cytoplasmic RNA in ribosome lysis buffer (20 mM Tris-HCl pH 7.4, 150 mM NaCl, 5 mM MgCl2, 1 mM DTT, 1% Triton X-100, Turbo DNase I 25U/ml). Extracted RNA was then digested by RNase I at room temperature for 45 minutes on a nutator. Digestion was stopped by adding SuperRNaseIn RNase inhibitor. The digested RNA was then loaded on ribosome buffer (lysis buffer without Triton-100 and DNase) with 2 ml of 1 M sucrose cushion and centrifuged at maximum speed on SW-32 rotor of Beckman Coulter ultracentrifuge at 4 degrees C for 7 hours. The supernatant was then poured out and the translucent pellet was then resuspended in 1 ml of Trizol reagent. RNA was extracted according to the Trizol reagent manufacturer’s instructions. Purified RNA was run in 15% polyacrylamide TBE-urea gel with synthesized 24 nt and 36 nt RNA oligos as markers for excision. Fragments between 24-36 nt were excised and extracted by freezing in dry ice bath with RNA extraction buffer (300 mM NaOAc pH 5.5, 1.0 mM EDTA, 0.25% SDS) for 30 minutes, followed by an overnight incubation at room temperature on a nutator. Diffused RNA in the RNA extraction buffer was then precipitated and purified by isopropanol. Purified RNA was then ligated with universal miRNA adaptor (NEB). Excessive adaptors were removed by running the ligation product in 15% polyacrylamide TBE-urea gel. Ligated RNA was then reverse transcribed by SuperScript III cDNA synthesis kit (Invitrogen). This first strand cDNA was purified from excessive RT primers through polyacrylamide gel electrophoresis and circularized by CircLigase. Circularized cDNA was PCR amplified to construct sequencing libraries and purified by polyacrylamide gel electrophoresis. Library was then quantified by KAPA library quantification kit and sequenced in Novaseq 6000 platform.

For analysis, raw FASTQ files were firstly trimmed by cutadapt using the universal miRNA adaptor sequence. Reads shorter than 20 bp after trimming were removed. Trimmed FASTQ files were firstly aligned against human rRNA index using bowtie aligner (Langmead, 2010).Unmatched FASTQ files were further aligned against the GRCh38 index using STAR (Dobin et al., 2013). Aligned bam files were analyzed by RiboCode software for 3-nt periodicity identification and a new GTF file containing actively translated ORFs was generated (Xiao et al., 2018), which was then used for count table generation by HTSeq software (Anders et al., 2014).

### Lipidomics analysis

Cells were FACS sorted directly into 100% isopropanol for lipid extraction. Analyses of polar and non-polar plasma lipids were conducted using an LC-MS system comprised of a Shimadzu Nexera X2 U-HPLC (Shimadzu Corp.) coupled to an Exactive Plus orbitrap mass spectrometer (Thermo Fisher Scientific). After centrifugation, to remove cellular debris, 10uL cell extracts in isopropanol were injected onto a 100 x 2.1 mm, 1.7 µm ACQUITY BEH C8 column (Waters). The column was eluted isocratically with 80% mobile phase A (95:5:0.1 vol/vol/vol 10mM ammonium acetate/methanol/formic acid) for 1 minute followed by a linear gradient to 80% mobile-phase B (99.9:0.1 vol/vol methanol/formic acid) over 2 minutes, a linear gradient to 100% mobile phase B over 7 minutes, then 3 minutes at 100% mobile-phase B. MS analyses were carried out using electrospray ionization in the positive ion mode using full scan analysis over 220–1100 m/z at 70,000 resolution and 3 Hz data acquisition rate. Other MS settings were: sheath gas 50, in source CID 5 eV, sweep gas 5, spray voltage 3 kV, capillary temperature 300°C, S-lens RF 60, heater temperature 300°C, microscans 1, automatic gain control target 1e6, and maximum ion time 100 ms. Raw data were processed using TraceFinder software (Thermo Fisher Scientific) for targeted peak integration and manual review of a subset of identified lipids and using Progenesis QI (Nonlinear Dynamics) for peak detection and integration of both lipids of known identity and unknowns. Lipid identities were determined based on comparison to reference plasma extracts and are denoted by total number of carbons in the lipid acyl chain(s) and total number of double bonds in the lipid acyl chain(s). Downstream statistical analysis and lipid class enrichment analysis were done by LipidSig software (Lin et al., 2021).

---

**Figure S1.**
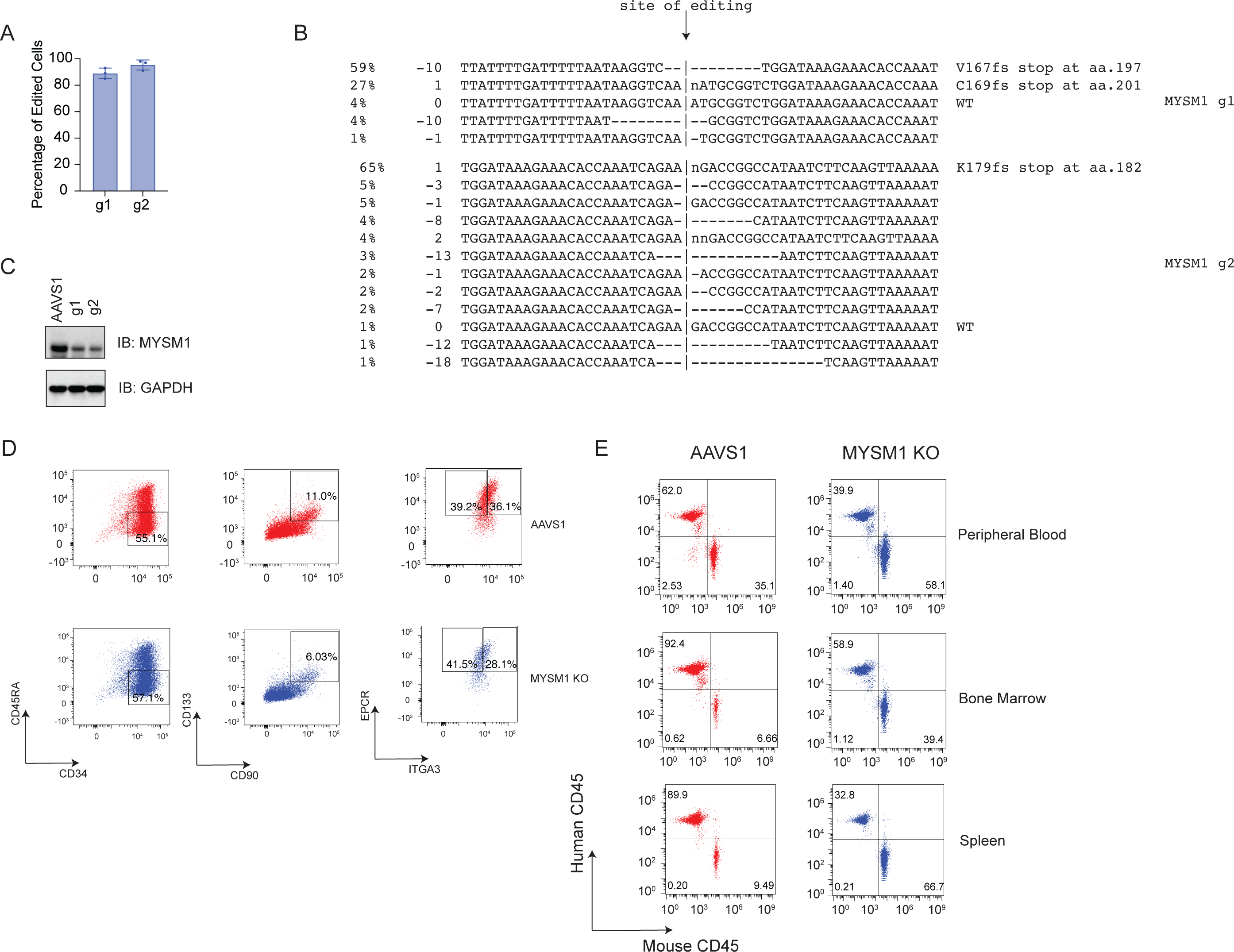
CRISPR editing analysis and flow cytometric analysis of HSC phenotyping, related to Figure 1. (A) ICE analysis of percentage of edited cells by two independent MYSM1 gRNAs. (B) Prediction of edited species by two MYSM1 gRNAs. Major edited species all result in stop codons at very early sites of the MYSM1 gene. (C) Western blot validation of successful knockout of MYSM1. (D) Representative flow cytometric analysis of HSC phenotyping of AAVS1 control or MYSM1 edited cells and gating strategy. (E) Representative flow cytometric analysis of human vs. mouse CD45^+^ cells after RBC depletion from indicated sites of NBSGW mice xenotransplanted with AAVS1 and MYSM1 edited cord blood CD34^+^ HSPCs.

**Figure S2.**
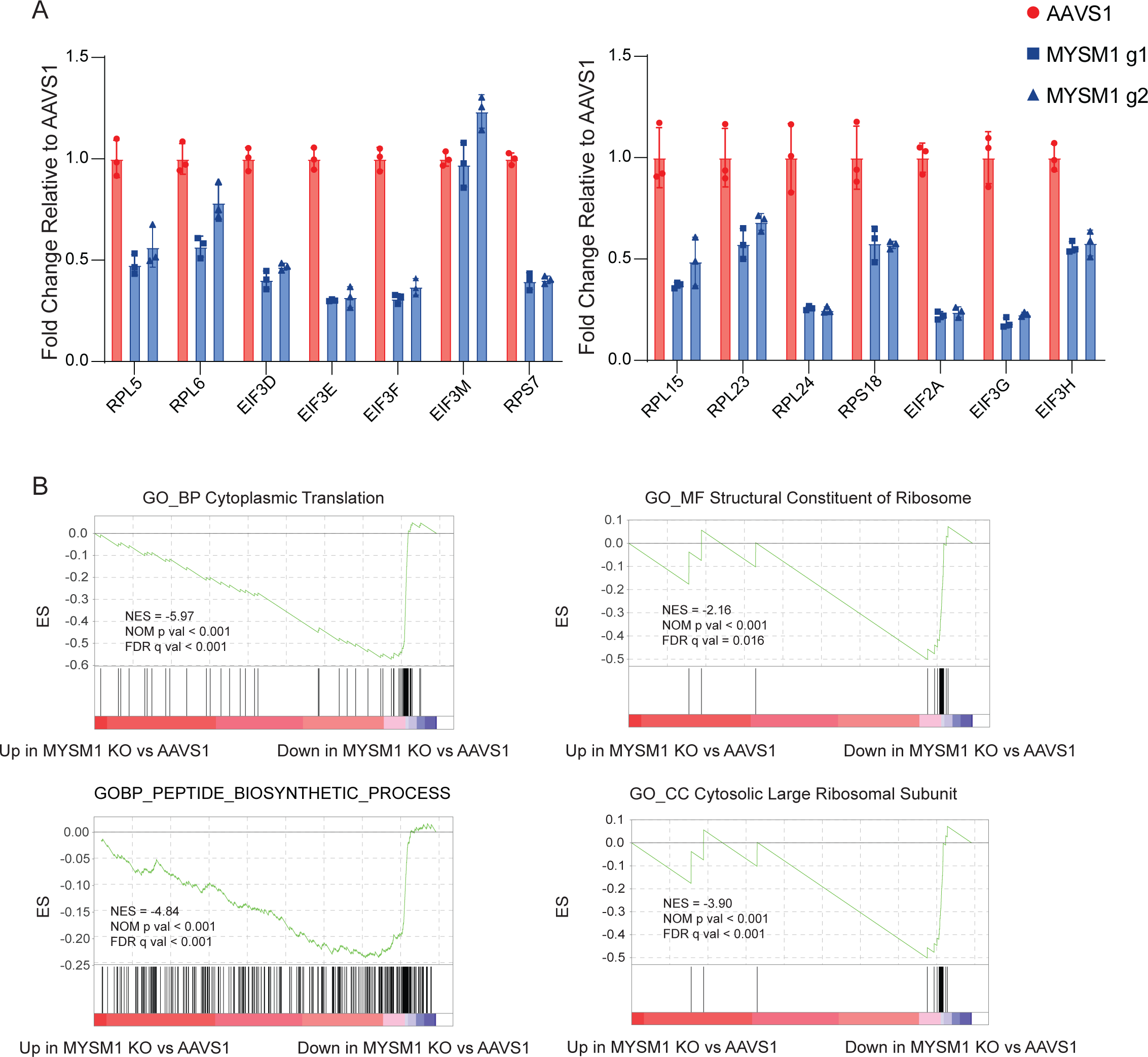
Transcriptional regulation of translation genes by MYSM1, related to Figure 2. (A) Additional gene set enrichment analysis (GSEA) showing ribosomal biogenesis and cellular translation are impaired with loss of MYSM1. (B) Real-time PCR validation of genes shown as downregulation due to loss of MYSM1 in scRNA-seq.

**Figure S3.**
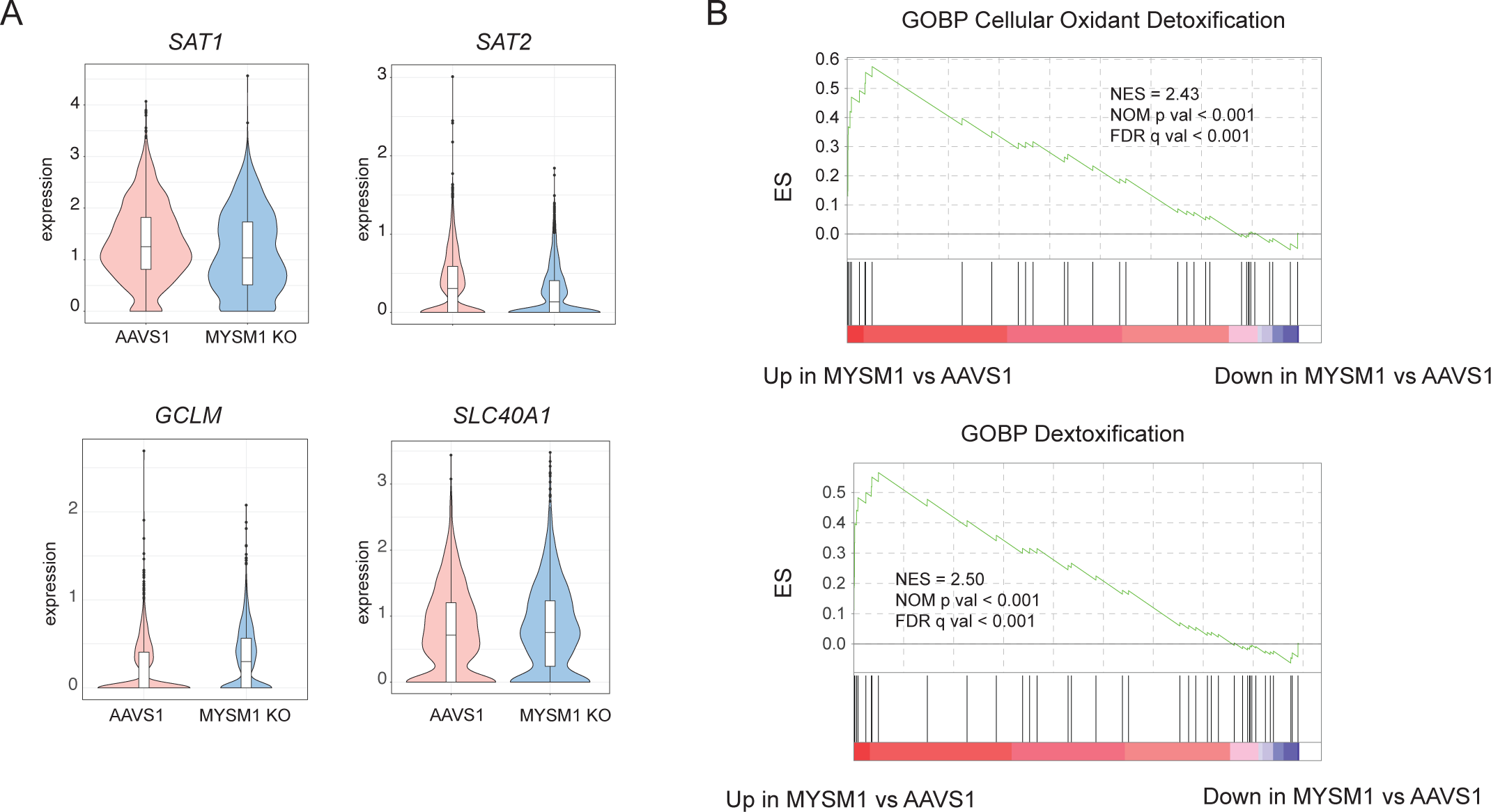
Upregulation of ferroptosis protective genes and downregulation of ferroptosis promoting genes upon loss of MYSM1, related to Figure 3. (A) Additional violin plot of ferroptosis gene expression in AAVS1 and MYSM1 KO HSCs. (B) Additional gene set enrichment analysis (GSEA) showing that cellular oxidant detoxification and general detoxification processes were significantly stimulated in MYSM1 KO HSCs.

**Figure S4.**
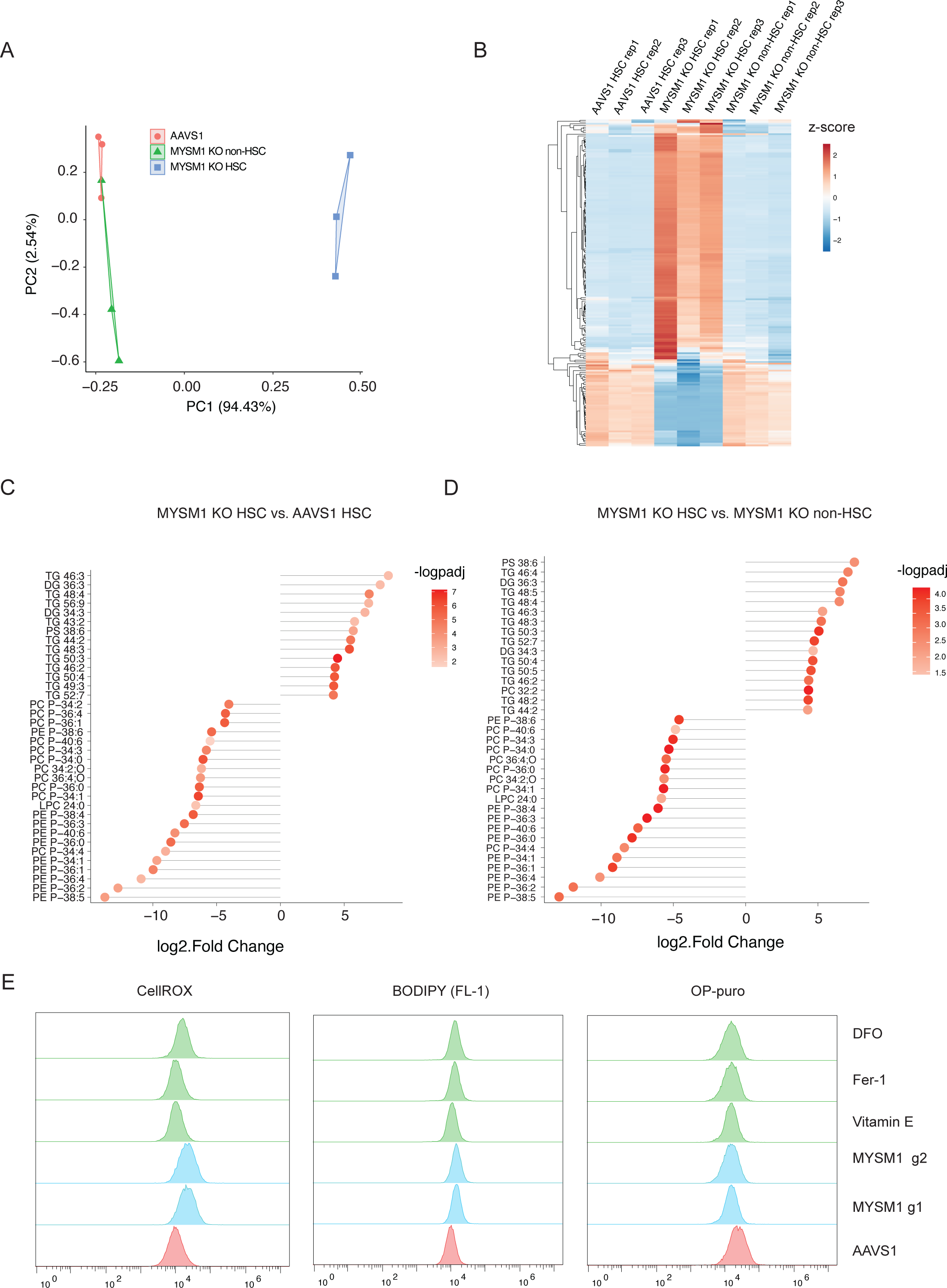
Increased oxidative stress and lipid peroxidation and depletion of membrane phospholipids upon loss of MYSM1, related to Figure 4. (A) Principal Component Analysis (PCA) of three replicates of the lipidomics of AAVS1 control, sorted CD90^+^CD133^+^ and CD90^-^ MYSM1 KO cells. (B) Unsupervised hierarchical analysis of three replicates of AAVS1 control, sorted CD90^+^CD133^+^ and CD90^-^ MYSM1 KO cells. (C) Bubble chart of selected enriched and depleted lipids of sorted CD90^+^CD133^+^ MYSM1 KO cells compared to AVVS1 control. (D) Bubble chart of selected enriched and depleted lipids of sorted CD90^+^CD133^+^ MYSM1 KO cells compared to CD90^-^ MYSM1 KO cells. (E) Representative flow cytometric histogram of CellROX, oxidized BODIPY and O-Propargyl-puromycin signals of sorted CD34^+^CD45RA^-^CD90^+^ cells with AAVS1 or MYSM1 editing, and MYSM1 edited cells with indicated chemical treatments.

**Figure S5.**
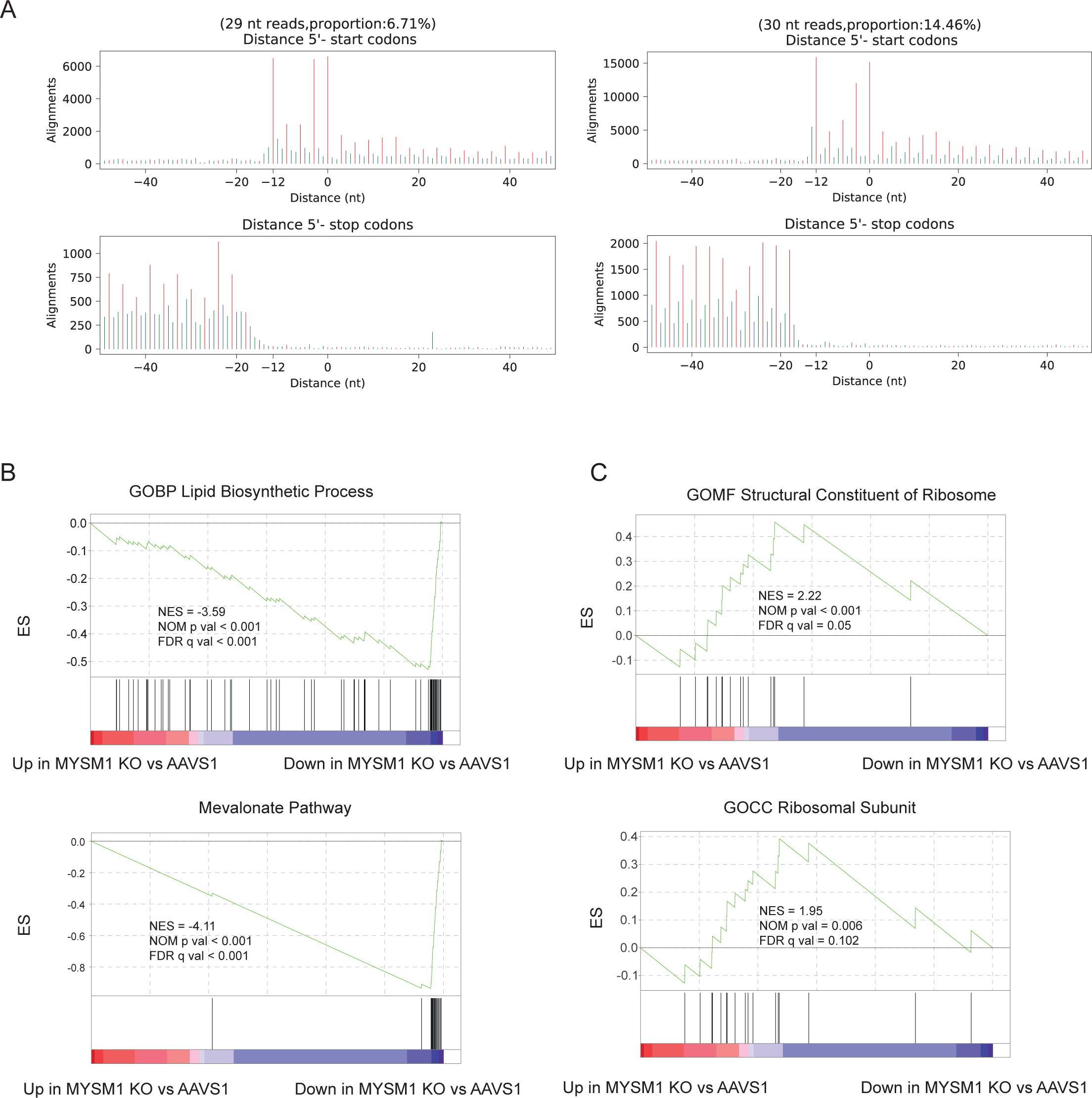
Quality control of ribo-seq experiment and GSEA of top genes whose translation is affected, related to Figure 5. (A) Identification of 3-nt periodicity of ribosome profiling sequencing (ribo-seq) data. (B) Additional gene set enrichment analysis (GSEA) showing that lipid synthesis related gene translation and mevalonate pathway translation were also significantly reduced in MYSM1 KO cells. (C) Additional gene set enrichment analysis (GSEA) showing that ribosome biogenesis and translation initiation gene translation was significantly increased.

**Figure S6.**
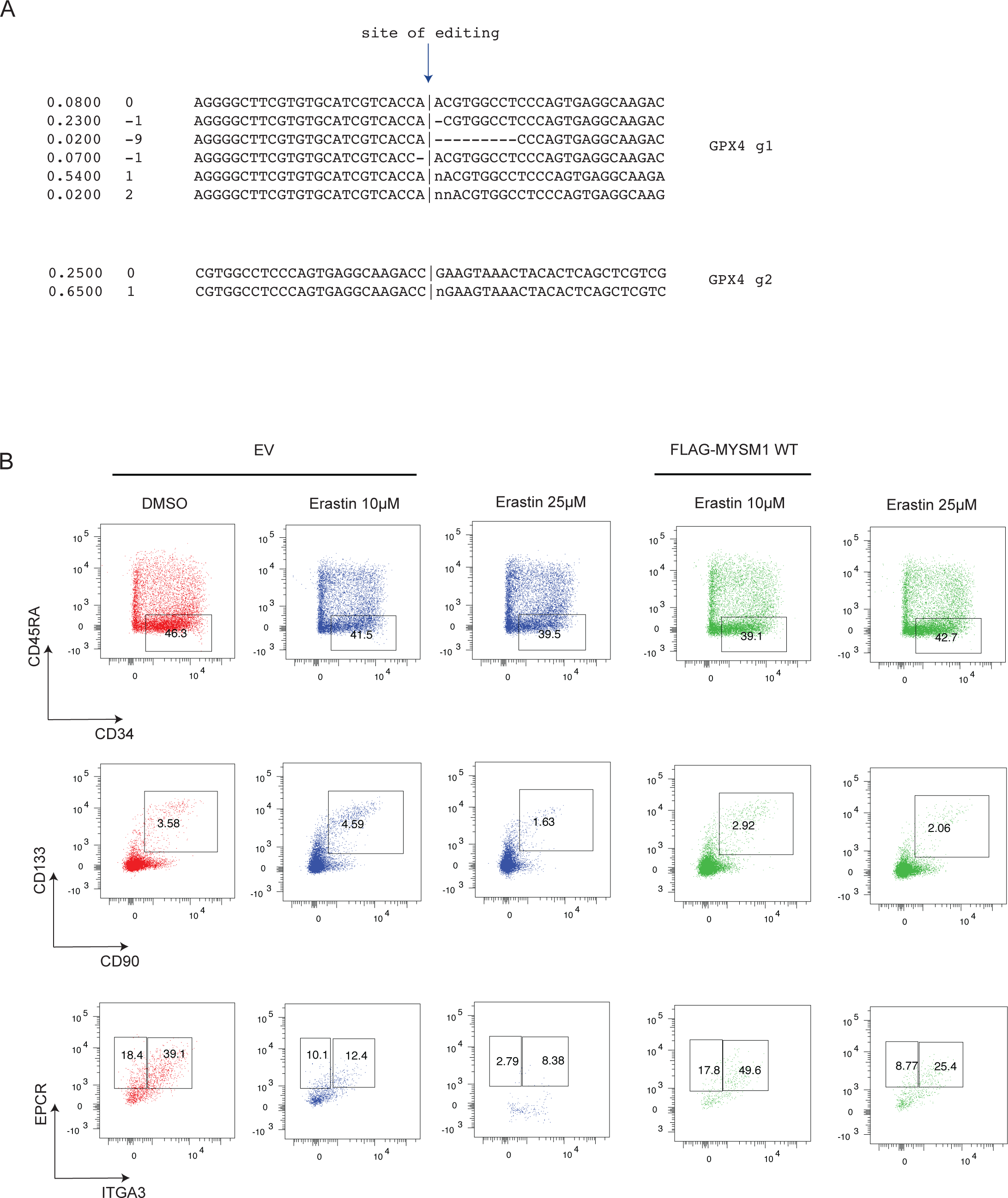
Flow cytometric analysis of HSC phenotyping upon ferroptosis induction, related to Figure 6. (A) Prediction of edit species by two independent gRNAs of GPX4. (B) Representative flow cytometric analysis of HSC phenotyping of CD34^+^ HSPCs treated with Erastin at indicated concentration transduced with empty vector (EV) or wildtype MYSM1.

## Supplementary Tables

**Table S1** Differentially expressed genes of MYSM1 KO vs AAVS1

**Table S2** GSEA analysis of differentially expressed genes of MYSM1 KO vs AAVS1

**Table S3** Lipidomics of MYSM1 KO HSC vs AAVS1 HSC

**Table S4** Lipidomics of MYSM1 KO non-HSC vs AAVS1 HSC

**Table S5** Lipidomics of MYSM1 KO HSC vs MYSM1 KO non-HSC

**Table S6** Differentially translated genes of MYSM1 KO vs AAVS1

**Table S7** GSEA of differentially translated genes of MYSM1 KO vs AAVS1

